# Production of SARS-CoV-2 virus-like particles in insect cells

**DOI:** 10.1101/2021.01.30.428979

**Authors:** Youjun Mi, Tao Xie, Bingdong Zhu, Jiying Tan, Xuefeng Li, Yanping Luo, Fei Li, HongXia Niu, Jiangyuan Han, Wei Lv, Juan Wang

## Abstract

Coronavirus disease (COVID-19) causes a serious threat to human health. To production of SARS-COV-2 virus-like particles (VLPs) in insect cells for vaccine development and scientific research. The E, M and S genes were cloned into multiple cloning sites of the new triple expression plasmid with one p10 promoter, two pPH promoters and three multiple cloning sites. The plasmid was transformed into DH10 Bac^TM^ *Escherichia coli* competent cells to obtain recombinant bacmid. Then the recombinant bacmid was transfected in ExpiSf9™ insect cells to generate recombinant baculovirus. After ExpiSf9™ infected with the recombinant baculovirus, the E, M, and S protein co-expressed in insect cells. Finally, SARS-CoV-2 VLPs were self-assembled in insect cells after infection. The morphology and the size of SARS-CoV-2 VLPs are similar to the native virions.

---

Coronavirus disease (COVID-19), caused by the severe acute respiratory syndrome coronavirus-2 (SARS-CoV-2), went on to ravage the world and caused the biggest pandemic 21st century. Compared with severe acute respiratory syndrome coronavirus (SARS-CoV) and Middle East respiratory coronavirus (MERS-CoV), severe acute respiratory syndrome coronavirus 2 (SARS-CoV-2) spreads faster with an infection index of about 2.6 [1, 2]. On October 15, 2020, more than 38.69 million people worldwide have been infected with the SARS-CoV-2, of which more than 1090,000 have died [3]. At present, there are no specific drugs for the coronavirus disease 2019 (COVID-19), many efforts have focused on neutralizing antibody and vaccine development [4]. Vaccines are the most economical and effective means to prevent and control infectious diseases [5]. There is an urgent need to develop SARS-CoV-2 vaccines to prevent and control the spread of the virus.

While many efforts are being made to design and develop vaccines for SARS-CoV-2. The World Health Organization estimates that there are about 133 vaccines under development [6]. Coronavirus vaccines generally fall into one of the following types: inactive or live-attenuated viruses, protein-based, virus-like particles (VLPs), viral vectors, and nucleic acid vaccines[7]. Inactivated viruses or live attenuated viruses require a large number of viruses to be cultured under biosafety level 3 (BSL3) conditions, and extensive safety testing is required, but the process is expensive, laborious and has a high safety risk. Protein-based subunit vaccines have poor immunogenicity due to incorrect folding of the target protein or poor display to the immune system, and require the addition of an adjuvant to induce a strong immune response. Nucleic acid vaccines cannot enter cells efficiently, need to be electroporated after injection, and mRNA is not very stable, multiple inoculations are required [8]. As a specific type of subunit vaccine, VLPs can mimic the natural morphology and structure of viruses, and have the characteristics of not containing viral genome nucleic acid, inability to replicate, and repeated vaccination of individuals. Because of VLPs strong immunogenicity, ability to elicit protective neutralizing antibodies and reliable safety, VLPs are strong candidates for vaccines design. At present, several vaccines based on VLPs are commercially available, these include human papillomavirus (HPV) vaccine and hepatitis B vaccine (HBV) [9]..

SARS-CoV, SARS-CoV-2 and MERS-CoV are all belong to the group of Betacoronaviruses (βCoVs) and have similar structures [10]. According to previous studies, the composition of SARS-CoV VLPs [6] and MERS-CoV VLPs [8] required the E, M, and S proteins co-expression in the cells. Similar to other βCoVs, The 3’ end of the SARS-CoV-2 genome encodes 4 main structural proteins, including the spike (S) protein, the envelope (E) protein, the membrane (M) protein and the nucleocapsid (N) protein [11]. We speculate that SARS-CoV-2 VLP also consists of the E, M and the S protein. In this study, a triple expression plasmid that co-expression of the E, M, S proteins was constructed. The Bac to Bac baculovirus insect expression system was used to achieve the co-expression of the E, M, and S proteins in ExpiSf9™ cells. Eventually, the E, M, S proteins self-assemble to form VLPs in the cell.

## MATERIALS AND METHODS

### Cell lines

ExpiSf9™ cells were presented by Mr. Ru Yi from the State Key Laboratory of Lanzhou Institute of Veterinary Medicine, Chinese Academy of Agricultural Sciences. Expi™Sf9 cells were maintained as suspension cultures in flasks with serum-free ExpiSf™ CD Medium (Gibco, USA) at 27°C with stirring at a speed of 125 rpm. Cell density was determined by microscopically counting the number of cells and cell viability was judged by trypan blue dye exclusion.

### Construction of the EMS triple expression plasmid

The dual expression plasmid pFastBacDual (Invitrogen, USA) was purchased from Thermo Fisher Scientific China Co., Ltd.; In order to obtain a single recombinant baculovirus that co-expressed the M, E and S proteins, a new triple expression vector was generated. In brief, a new SV40 poly A tail, a new pPH promoter and a *NdeI* restriction site were inserted into the *EcoRI* and *HindIII* restriction sites after pPH promoter of pFastBacDual. The new triple expression vector has one p10 promoter and two pPH promoters. For co-expression of the E, M, and S proteins, the codon optimized E, M, S genes of SARS-CoV-2 (GenBank accession No. MN908947.3) cloned into the triple expression vector. The E gene cloned into the double *KpnI* and *XhoI* restriction sites under the control of the p10 promoter, the M gene inserted into the *BamHI* and *EcoRI* restriction sites under the control of the pPH promoter, and the S genes cloned into the double *NdeI* and *HindIII* restriction sites under the control of other pPH promoter. Finally, the triple expression plasmid named EMS was generated and verified by DNA sequencing (BGI, China).

### Generation of recombinant baculovirus

Recombinant baculovirus were generated by using a Bac-to-Bac expression system (Invitrogen, USA) according to the manufacturer’s instructions. Briefly, the EMS plasmid was transformed into Top10 competent (Transgen, China). After re-extracting the EMS plasmid, it was transformed into DH10 Bac^TM^ *E. coli* competent (Invitrogen, USA). White colonies were screened in LB media containing the antibiotics gentamicin (7μg/ml), tetracycline (10 μg/ml), kanamycin (50 μg/ml) and X-Gal (5-bromo-4-chloro-3-indolyl-β-D-galactopyranoside) and IPTG (isopropyl-β-D-thiogalactopyranoside). After twice cycles of white colony screening, recombinant Bacmid DNA were isolated and transfected into ExpiSf9^TM^ cells. Transfected ExpiSf9™ cells cultured at 27°C and 125rpm, until the cytopathic rate reached 30%, the cell supernatant was collected to obtain the recombinant baculovirus. Using PCR to verify the genes of interest in the recombinant bacmid. The primers for PCR are shown in Table 1.

**Table 1.**
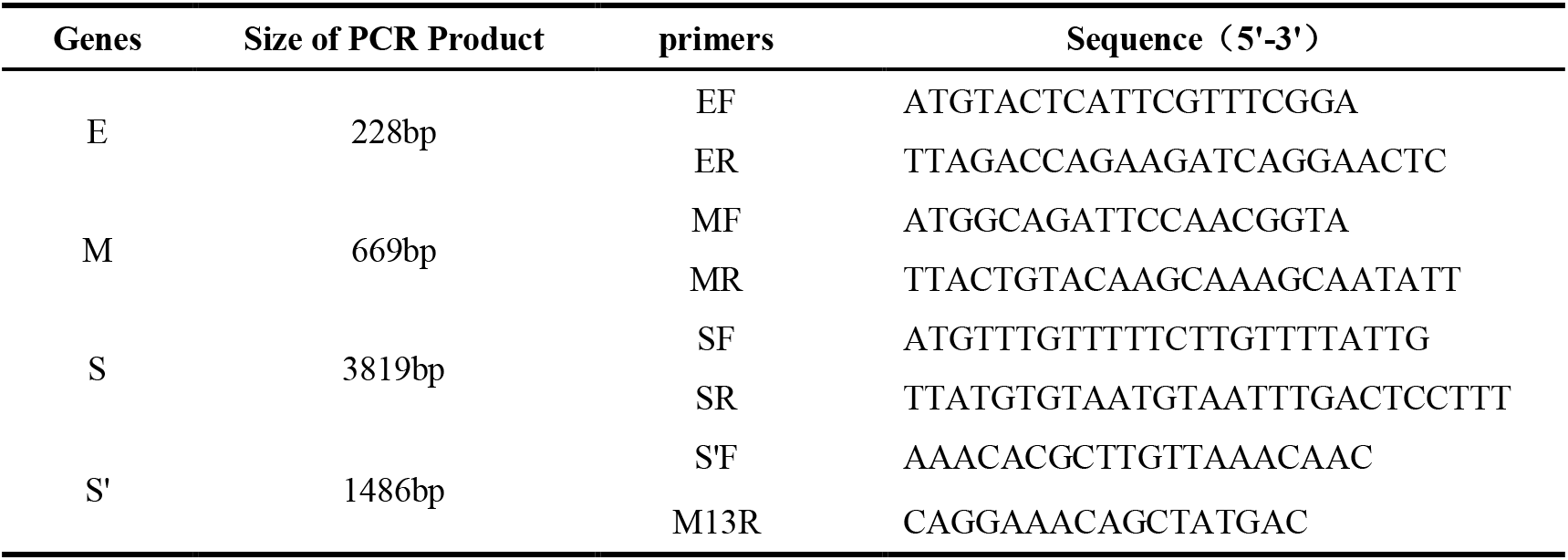
Analyze recombinant bacmid DNA by PCR

### Preparation of EMS VLPs

Incubating ExpiSf9™ cells for 18 hours, add ExpiSf™ Enhancer (Invitrogen, USA) to the cell culture. 24 hours after ExpiSf™ Enhancer addition, infect cells with baculovirus in a volume of 50:1. Cells were collected on day 4 by centrifugation at 3000 *g* for 5 minutes, freeze-thaw repeatedly for 3 times, and discard the pellet after centrifugation at 8000 *g* for 30 minutes. The supernatant was ultra-centrifuged at 100000 *g* for 1 hour at 4°C, and the pellets were resuspended in phosphate-buffered saline (PBS) at 4°C overnight. EMS VLPs were through a 30%-40%-50% discontinuous sucrose gradient at 100,000 *g* for 2 hours at 4°C. The white bands between 30-40% were collected and diluted with PBS and pelleted at 100000 *g* for 1 hour at 4°C. The VLPs was collected and resuspended in PBS overnight at 4°C. EMS VLPs were store −80°C for the following analysis.

### Western Blot Analysis

The VLPs were characterized by Western blot analysis and electric microscopic observation. For Western blot analysis, VLPs sample were subjected to SDS-PAGE using a 10% gel, followed by transfer to poly vinylidene difluoride (PVDF) membranes. The PVDF membranes were then blocked with TBST (10 mM Tris-HCl, 150 mM NaCl, 0.5% Tween 20) containing 5% skim-milk powder. The S protein of VLP was detected by Western blotting, using an anti-SARS-CoV-2 S polyclonal rabbit antibody (Sino Biological, China). The expression of the E and the M protein were also verified by polyclonal antibodies (data not shown). Alkaline phosphatase-conjugated goat-anti-rabbit IgG (immunoway, China) were used as the secondary antibodies to label the protein bands.

### Electron microscopy

For transmission electron microscopy. 5ul VLPs samples were applied onto a carbon-coated film. 2 minutes later, the samples was removed with filter paper. Then, 8 μl of 1% phosphotungstic acid was applied onto the grid, and the samples were stained for 60s. The staining solution was removed with filter paper, and the grid was dried for 30 min at room temperature. After being stained, the sample was observed using a FEI Talos F200C transmission electron microscope (FEI, Czech Republic) at 200 kV and 100-200 k-fold magnification.

## Results

### Construction of EMS triple expression plasmid

To co-expression the SARS-CoV-2 E, M, S proteins in insect cells, a new triple expression vector was constructed. The triple expression vector contains one pP10 promoter, two pPH promoters and three multiple cloning sites (Fig1 b). Then the SARS-CoV-2 E, M and S genes were cloned into multiple cloning sites respectively to generate EMS plamsid (Fig1 c).

**Fig 1.**
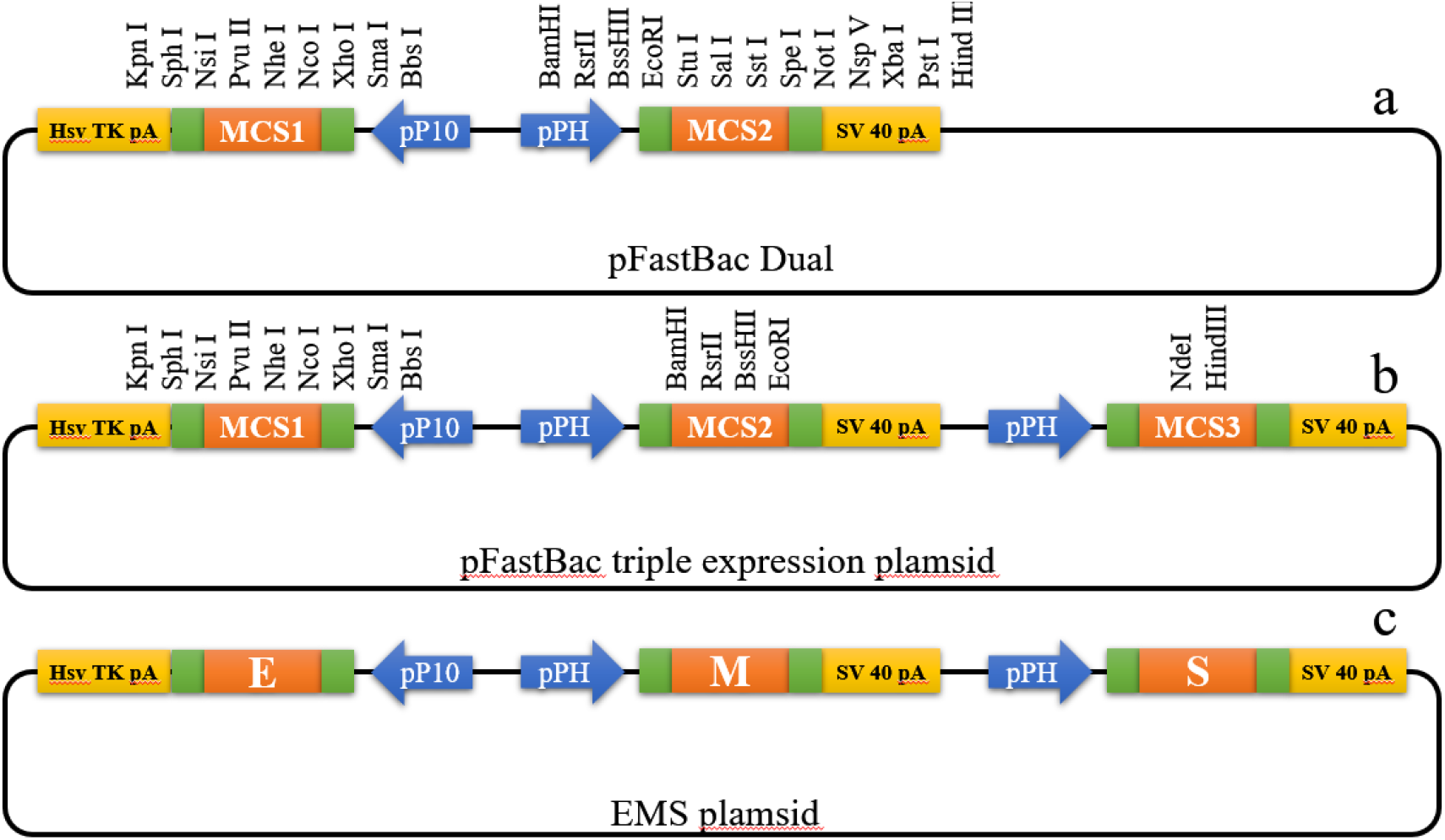
Construction of EMS plasmid a pFastBac Dual b pFastBac triple expression plasmid c EMS plasmid

### Generation of recombinant bacmid DNA

The EMS **r**ecombinant plasmid was transformed into DH10 Bac^TM^ *E. coli* competent Cells. Blue and white colonies were visible on the LB screening plate after 48 hours (Fig. 2a). Pick white colonies for the second screening to obtain positive colonies which containing the recombinant bacmid (Fig. 2b). To verify the E, M, and S genes in the recombinant bacmid by using PCR analysis. The target bands can be obtained at the corresponding positions (Fig. 3)

**Fig 2.**
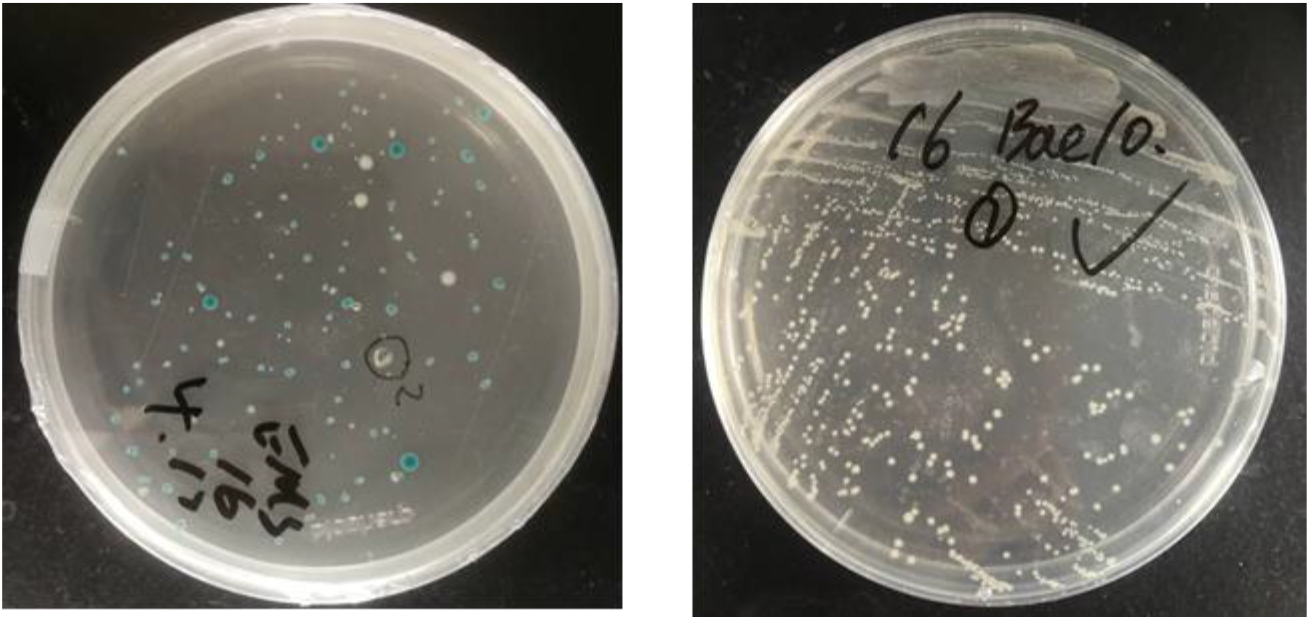
Transformation and secondary screening a The EMS vector was transformed into DH10 Bac^TM^ competent cells b Secondary screening after transformation

**Fig 3.**
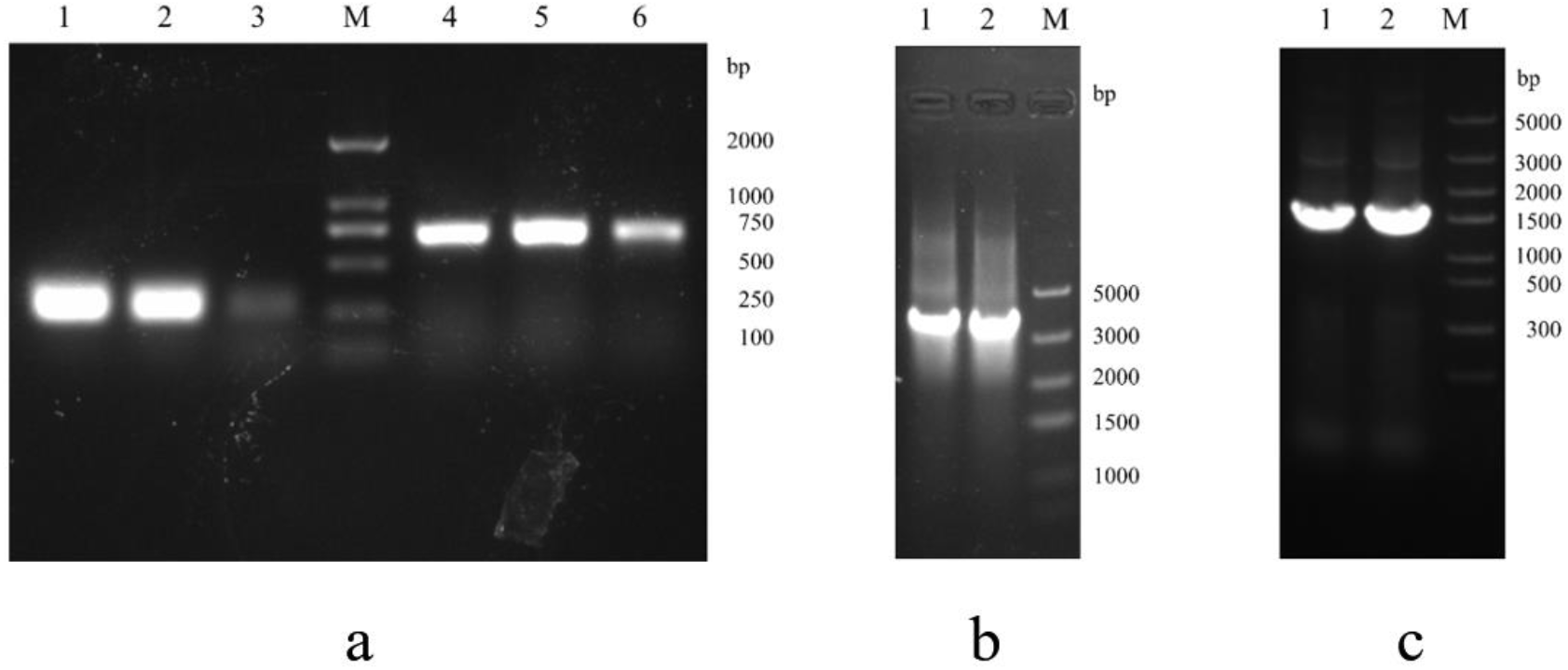
Verify the phenotype a 1,2,3 Identification of the E gene by PCR;4,5,6 Identification of the M gene by PCR b 1,2 Identification of the S gene by PCR c 1,2 Identification of S’ DNA fragment by PCR

### Recombinant baculovirus production

For producing of recombinant baculovirus, isolating bacmid DNA to transfect ExpiSf9™ insect cells. After the baculovirus is added to the cells, observation the signs of infection. The swollen cells with enlarged nuclei indicate that the cell is infected by baculovirus (Fig. 4). After trypan blue staining, the cell death rate was increased. Collecting the recombinant baculovirus from the cell culture medium when the cells appear characteristics typical of late to very late infection.

**Fig 4.**
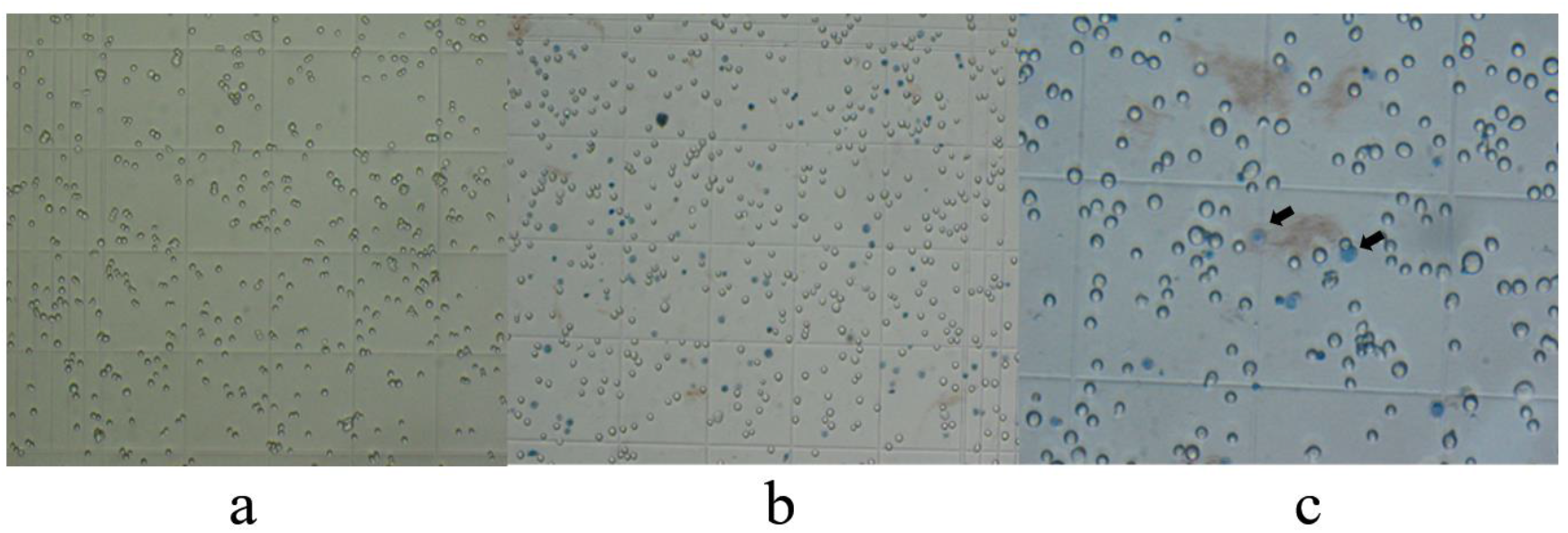
Cytopathic signs observed in ExpiSf9^TM^ cells a ExpiSf9^TM^ cells (×10) b Transfected ExpiSf9^TM^ cells (×10) c Transfected ExpiSf9^TM^ cells (× 40)

### Production and characterization of SARS-CoV-2 VLPs

To product SARS-CoV-2 VLPs in insect cells, Sf9 cells were infected with recombinant baculovirus. 96 hours after infection, the cells were harvested and SARS-CoV-2 VLPs were purified by sucrose gradient centrifugation. To confirm the S protein was incorporated within the VLPs, purified VLPs were analyzed by Western blot using SARS-CoV-2 S protein polyclonal antibody. The result showed that VLPs contained SARS-CoV-2 S proteins (Fig 5). The morphology of SARS-CoV-2 VLPs was investigated by electron microscopy. The SARS-CoV-2 VLPs exhibited spheriform structures, and the diameters of VLPs were approximately 100 nm (Fig 6). These results indicated that SARS-CoV-2 VLPs autonomously assemble in insect cells infected with recombinant baculovirus, and are structurally similar to the native virions[12].

**Fig.5.**
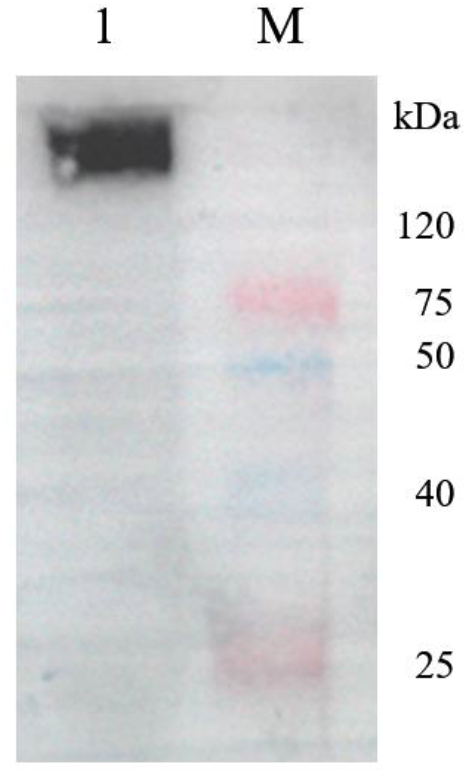
Western-blotting identification of SARS-CoV-2 S protein expressed on VLPs 1 identification of SARS-CoV-2 S protein with S antibody M Protein maker

**Fig 6.**
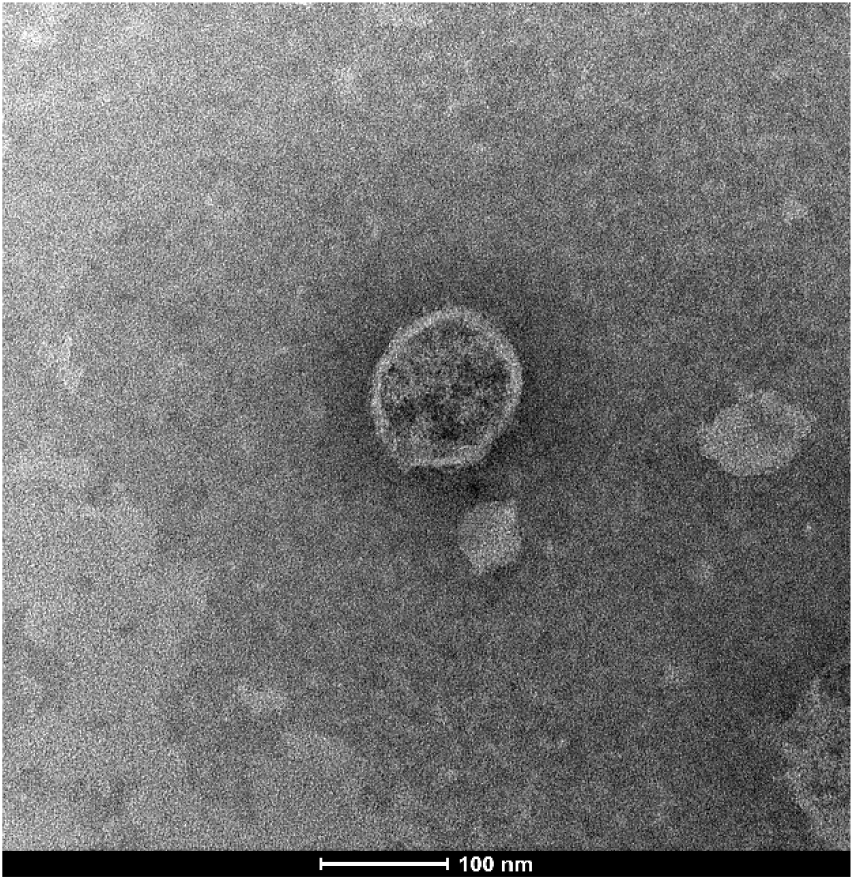
VLP observed under electron microscope

## CONCLUSION

Although some countries have successfully controlled the COVID-19 epidemic, on the face of ongoing death and destruction caused by SARS-CoV-2 virus, the urgent need for developing vaccines to against the spread of the virus. At present, SARS-CoV-2 related traditional vaccines and new generation vaccines are under development, but there is no effective vaccine approved for use. Because of their unique advantages, VLPs have been use as vaccine platform. Baculovirus system is especially suitable for virus-like particle production, since this system allows more than one gene to be co-expressed in a cell.

SARS-CoV-2, SARS-Cov, and MERS-CoV are all belong to β-CoV, have similar virus structures which consist of 4 structural proteins N, E, M, S[11]. Eduardo Mortola reported that the M, E and S proteins self-assembled to form SARS-CoV VLPs when co-transfection of the E, M, and S baculovirus in Sf9 insect cells[13]. Xinya Lu confirmed that SARS-Cov VLPs are immunogenic and can cause strong SARS-Cov VLPs specific humoral and cellular immune responses in mice [14]. Similarly, the co-infection of MERS CoV E, M and S recombinant baculoviruses in insect cells produces VLPs with similar morphological signature to the native virions. MERS-CoV VLPs are immunogenic, able eliciting robust levels of specific humoral and cell-mediated immunity in rhesus monkeys. Rhesus monkeys vaccinated MERS-CoV VLPs with alum adjuvant induced high titer virus neutralizing antibodies and triggered T helper 1 cells (Th1) mediated immunity[15].

In this study, a new triple expression vector was constructed, then the E, M, and S gene were cloned into the triple expression vector to generate EMS plasmid. The EMS vector was transformed into DH10 Bac^TM^ competent cells, and the recombinant bacmid was obtained after twice screening. Then the recombinant bacmid was transfected in ExpiSf9™ insect cells to obtain recombinant baculovirus. After ExpiSf9™ infected with the recombinant baculovirus, the E, M, and S protein co-expressed in cells to form VLPs by self-assembly. To our knowledge, a successful construction of SARS-CoV-2 VLPs via insect expression system has not yet been reported. In the future, humoral and cellular immunogenicity of SARS-CoV-2 VLPs in animal models will be further evaluated. In addition, VLPs can also be used to study the pathogenesis of COVID-19.

## ACKNOWLEDGEMENTS

This study was Supported by the Fundamental Research Funds for the Central Universities (lzujbky-2020-sp06 and lzujbky-2017-22), supported by the Key research and development program of Gansu Province in 2020 (20YF8FA072) and by Lanzhou science and technology planning project(2020-XG-33).

## Notes

### Competing Interest Statement

The authors have declared no competing interest.

